# Transcriptomically-inferred PI3K activity and stemness show a counterintuitive correlation with *PIK3CA* genotype in breast cancer

**DOI:** 10.1101/2020.07.09.195974

**Authors:** Ralitsa R. Madsen, Oscar M. Rueda, Xavier Robin, Carlos Caldas, Robert K. Semple, Bart Vanhaesebroeck

## Abstract

A PI3Kα-selective inhibitor has recently been approved for use in breast tumours harbouring mutations in *PIK3CA*, the gene encoding PI3Kα. Preclinical studies have suggested that the PI3K/AKT/mTORC1 signalling pathway influences stemness, a dedifferentiation-related cellular phenotype associated with aggressive cancer. No direct evidence for such a correlation has been demonstrated to date in human tumours. In two independent human breast cancer cohorts, encompassing nearly 3,000 tumour samples, transcriptional footprint-based analysis uncovered a positive linear association between transcriptionally-inferred PI3K signalling scores and stemness scores. Unexpectedly, stratification of tumours according to *PIK3CA* genotype revealed a “biphasic” relationship of mutant *PIK3CA* allele dosage with these scores. Relative to tumour samples without *PIK3CA* mutations, the presence of a single copy of a hotspot *PIK3CA* variant was associated with lower PI3K signalling and stemness scores, whereas tumours with multiple copies of *PIK3CA* hotspot mutations showed higher PI3K signalling and stemness scores. This observation was recapitulated in a human cell model of heterozygous and homozygous *PIK3CA^H1047R^* expression. Collectively, our analysis provides evidence for a signalling strength-dependent PI3K-stemness relationship in human breast cancer, which may aid future patient stratification for PI3K-targeted therapies.

## INTRODUCTION

Activating mutations in *PIK3CA* are among the most common somatic point mutations in cancer, together with inactivation or loss of the tumour suppressor PTEN, a negative regulator of class I phosphoinoside 3-kinase (PI3K) enzymes [1–3]. PI3Kα-selective inhibitors are now making good progress in the clinic [4], with the PI3Kα-specific inhibitor alpelisib (Piqray/NVP-BYL719; Novartis) received approval for the treatment of advanced hormone-receptor (HR)-positive, HER2-negative breast cancers, following a randomised phase III trial evaluating alpelisib with the oestrogen receptor (ER) antagonist fulvestrant *versus* fulvestrant alone [5]. The trial concluded that a clinically-relevant benefit of the combination therapy was more likely in patients with *PIK3CA*-mutant tumours [5]. The FDA approval of alpelisib was accompanied by approval of the companion diagnostic therascreen^®^ *PIK3CA* test (QIAGEN) which detects 11 *PIK3CA* hotspot mutations. Despite these advances, a substantial proportion of patients with *PIK3CA*-mutant tumours failed to improve on the combination therapy [5], highlighting the need for further refinement of current patient stratification strategies.

Experimental evidence suggests that heterozygous expression of a strongly activating *PIK3CA* mutation alone is insufficient to transform cells *in vitro* or to induce tumourigenesis *in vivo* (reviewed in Ref. [6]). This is supported by observations of people with disorders in the *PIK3CA*-related overgrowth spectrum (PROS) which is caused by the same spectrum of *PIK3CA* mutations found in cancer, but does not feature discernible excess risk of adult malignancy [6]. It thus appears that additional events are required for cell transformation, possibly in the PI3K pathway itself. In this regard, we and others have recently shown that many *PIK3CA*-associated cancers harbour multiple independent mutations activating the PI3K pathway, including multiple *PIK3CA* mutations in *cis* or *trans* [3,7–10].

Overexpression of wild-type *Pik3ca* or the hotspot *Pik3ca^H1047R^* mutation has been linked to dedifferentiation and stemness in murine models of cancer [11–17], particularly of the breast, but *Pik3ca* gene dose-dependent regulation has not been addressed. Pluripotent stem cells (PSCs) share key characteristics with cancer cells, including developmental plasticity, the capacity for indefinite self-renewal, rapid proliferation and high glycolytic flux [18]. We recently reported that human PSCs with two endogenous alleles of the strongly activating cancer hotspot mutation *PIK3CA^H1047R^* exhibit pronounced phenotypic differences compared to isogenic cells heterozygous for the same *PIK3CA* variant [8]. These differences include partial loss of epithelial morphology, widespread transcriptional reprogramming and self-sustained stemness *in vitro* and *in vivo* [8], none of which were observed in heterozygous *PIK3CA^H1047R^* cells. Collectively these findings emphasise the importance of *PIK3CA* mutation dose, and its inferred functional correlate, PI3K signalling strength, in determining the cellular consequences of mutational activation of this pathway.

Stemness or dedifferentiation, accompanied by re-expression of embryonic genes, is a feature of aggressive tumours [19,20]. Beyond direct histopathological analyses, this has been supported by computational analyses examining a tumour’s expression of defined PSC gene signatures [19–22]. With the continuing collection and curation of multi-omics datasets by the cancer community, such signatures can now be employed *en masse* to study how cancer-specific stemness relates to other biological processes of interest. This can, however, be challenging for highly dynamic processes such as signalling pathway activity which is best inferred using temporal protein-based measurements. Such measurements are not available for most human tissue samples. A complementary approach is the use of transcriptional “footprints” of pathway activation, derived from the systematic curation of gene expression data obtained from direct perturbation experiments [23–25]. Given the slower time scale of gene expression regulation relative to acute signalling changes at the protein level, transcriptional footprint analyses can be thought of as providing an integrated measure of pathway activity over a longer time scale. The power of such analyses has been best demonstrated by The Connectivity Map Resource, which enables the discoveries of gene and drug mechanisms of action on the basis of common gene-expression signatures [26,27].

Here, we set out to determine whether a signalling strength-dependent PI3K-stemness link exists in human breast cancer, and to provide a systematic characterisation of relevant clinical and biological correlates. We used established, open-source methods to infer PI3K signalling activity and stemness scores from publicly available transcriptomic data from nearly 3,000 primary human breast tumours. Our analyses reveal a positive, linear relationship between PI3K signalling and stemness scores, and uncover a surprising and unanticipated ‘biphasic’ relationship between these scores and mutant *PIK3CA* allele dosage. This suggests a potential utility for combined functional genomics and genotype assessments in future patient stratification for PI3K-targeted therapy. Consistent with prior cell biology studies, breast tumour transcriptomic analyses revealed strong clustering of PI3K and stemness scores with MYC-related biological processes, including proliferation and glycolysis. With the advent of routine tumour gene expression analyses, further dissection of the mechanisms driving these associations may enable much-needed further therapeutic advances.

## RESULTS

### Transcriptional indices of PI3K pathway activity in breast cancer are positively associated with stemness and tumour grade

The molecular features of stemness can be captured by gene signatures derived by computational comparisons of pluripotent stem cells and differentiated derivatives. Among the first such signatures was PluriNet (n = 299 genes; **Supplementary Table 1**), generated with machine learning methods [28], and applied below to primary breast cancer samples. To evaluate PI3K pathway activity in the same samples, we used the “HALLMARK_PI3K_AKT_MTOR_SIGNALING” gene set from the Broad Institute Molecular Signature Database (MSigDB). This gene set consists of 105 genes upregulated upon PI3K pathway activation across multiple studies [24] (**Supplementary Table 2**), thus corresponding to a gene expression footprint of PI3K pathway activation. Of note, only 4 genes were shared between the PI3K activity and stemness gene lists, precluding a direct confounding effect on the relationship between stemness and PI3K activity scores tested here.

We used Gene Set Variation Analysis (GSVA) [29], an open-source method, to calculate stemness and PI3K activity scores on the basis of the aforementioned gene expression signatures, independently in breast cancer tumours with available transcriptomic data from the METABRIC (*n* = 1980; used for primary analyses) and TCGA patient cohorts (n = 928; used for secondary analyses). The PI3K activity score in METABRIC breast tumours correlated significantly with the stemness score (**Fig. 1A**; Spearman’s Rho = 0.49, p<2.2e-16) as well as tumour grade status (**Fig. 1B**), a measure of tumour dedifferentiation based on histopathological assessment. A similar linear relationship between PI3K activity and stemness scores was also found in TCGA breast cancers (**Fig. 1C**; Spearman’s Rho = 0.42; p<2.2e-16).

**Fig. 1.**
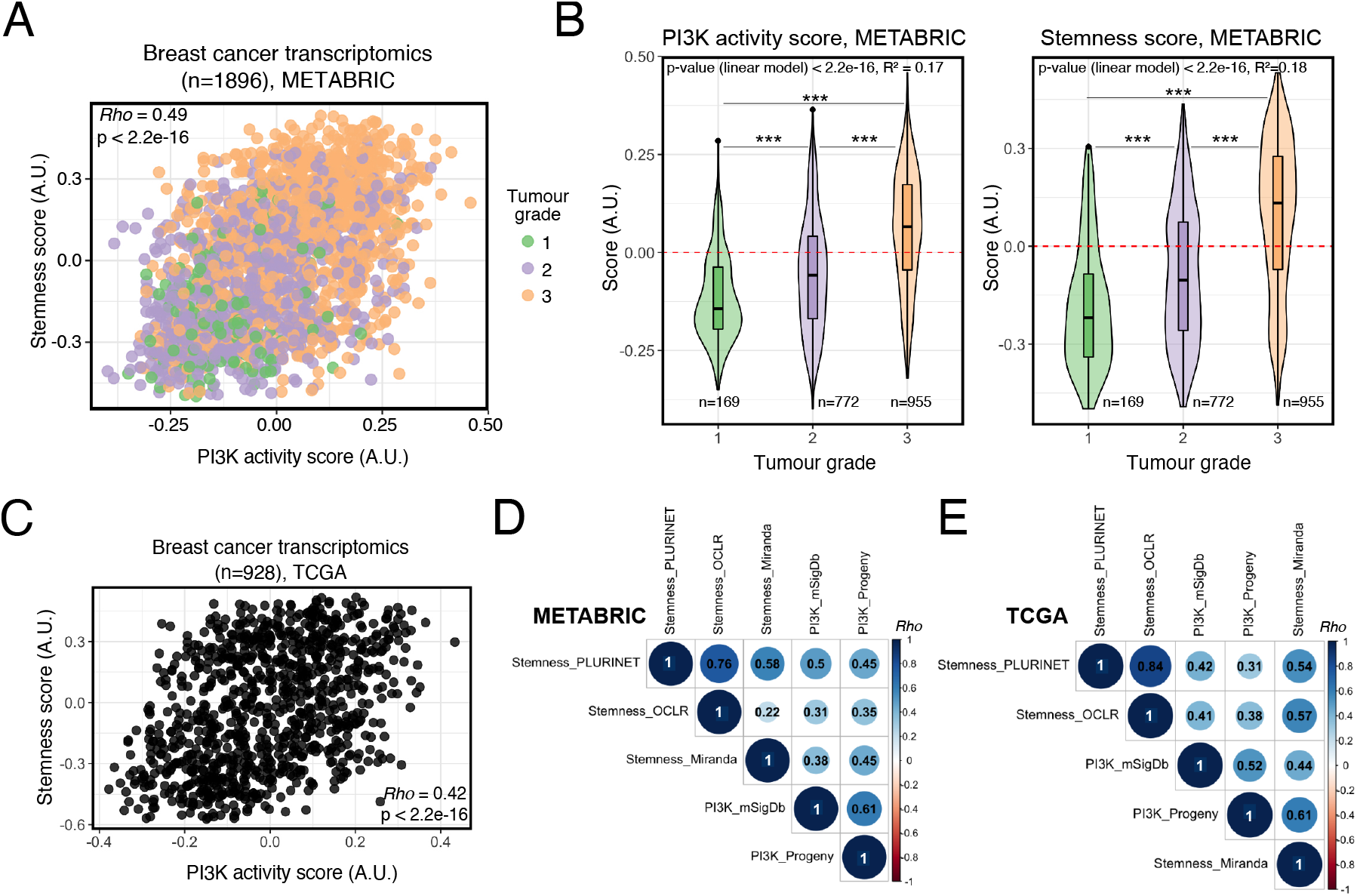
Strong positive association between transcriptionally-inferred PI3K pathway activation and breast tumour stemness/grade. **(A)** Rank-based (Spearman’s *Rho*) correlation analysis of the relationship between transcriptionally-inferred PI3K activity and stemness scores, evaluated across METABRIC breast cancer transcriptomes. Scores were determined using Gene Set Variation Analysis (GSVA) with mSigDb “HALLMARK_PI3K_AKT_MTOR_SIGNALING” (for PI3K activity score) and “MUELLER_PLURINET” (for stemness score) gene signatures [24,28,29]. Gene lists used are included in Supplementary Tables 1 and 2. **(B)** PI3K activity and stemness score distributions across breast cancer grade (METABRIC). *** p < 0.001 according to one-way ANOVA with Tukey’s Honest Significant Differences method. The global p-value for each linear model is indicated within each plot. **(C)** As in (A) but based on TCGA breast invasive carcinoma (BRCA) transcriptomic data. **(D)** Rank-based correlation analyses of the stemness (PluriNet-based) and PI3K (mSigDb-based) scores used in the current study against published and independently-derived transcriptional indices for stemness and PI3K signalling, across METABRIC breast cancer transcriptomic data. Individual *Rho* coefficients are shown within the respective circles whose sizes are matched accordingly. Only significant correlations are shown (family-wise error rate < 0.05). **(E)** As in (D) but based on TCGA BRCA transcriptomic data. The Stemness_OCLR score is based on a machine-learning-derived stemness signature [21]; the Stemness_Miranda score is based on a modification of the stemness signature of Palmer et al. [20,22]. The PI3K_Progeny score is based on the analysis of benchmarked pathway-responsive genes as described in Ref. [23].

To ascertain the ability of our approach to capture *bona fide* features of stemness and PI3K signalling from transcriptomic data, we next performed pairwise-correlations with independently-derived transcriptomic indices for each phenotype. Across both METABRIC (**Fig. 1D**) and TCGA (**Fig. 1E**) breast tumours, the PluriNet-derived stemness score showed good concordance with alternative stemness scores obtained using Malta *et al*.’s one-class logistic regression (OCLR)-based signature [21], or the signature from Miranda *et al*. [22], a modified version of a gene set initially developed by Palmer *et al*. [20]. The strongest correlations (Spearman’s Rho > 0.7) were between PluriNet and the OCLR-based signature, both of which were derived using distinct machine learning algorithms.

To strengthen our observations, we next applied PROGENy to obtain an independent measure of PI3K activity on the basis of the transcriptomic footprint. Instead of the enrichment score calculated by GSVA, PROGENy uses a linear model to infer pathway activity from the expression of 100 pathway-responsive genes [23]. The GSVA- and PROGENy-derived PI3K scores exhibited a significant positive correlation (Spearman’s Rho > 0.5) across both METABRIC and TCGA breast cancers (**Fig. 1D, 1E**). Consistently, the two PI3K activity scores also exhibited similar positive correlations with all three stemness indices (**Fig. 1D, 1E**).

Taken together, these results provide evidence for the existence of a positive relationship between stemness and overall PI3K activity in human breast cancer.

### Stemness and PI3K activity scores differ across breast cancer tumour subtypes

Using our GSVA-based stemness and PI3K activity scores, we next sought to determine their relationship with clinical breast cancer subtype. Upon stratification of METABRIC breast cancers into those with “high” and “low” PI3K activity scores, we found that around 45% of tumours with a high PI3K activity score were ER-negative, in contrast to 4% of tumours with low PI3K activity scores (**Fig. 2A**). In TCGA, the corresponding percentages were 33% and 7% (**Fig. S1A**). Consistently, PI3K activity and stemness scores were highest in the more aggressive PAM50 breast cancer subtypes (**Fig. 2B**), including Basal, HER2 and Luminal B. These findings are in line with independent studies relying on alternative indices and methods for quantifying PI3K signalling and stemness in separate analyses [19,21,30–32]. Importantly, the correlation of a high PI3K activity score with ER-negativity contrasts with the known enrichment of *PIK3CA* mutations in ER-positive breast tumours [32,33], which were also reproduced by our analyses (**Fig. 2C**, **Fig. 2D**).

**Fig. 2.**
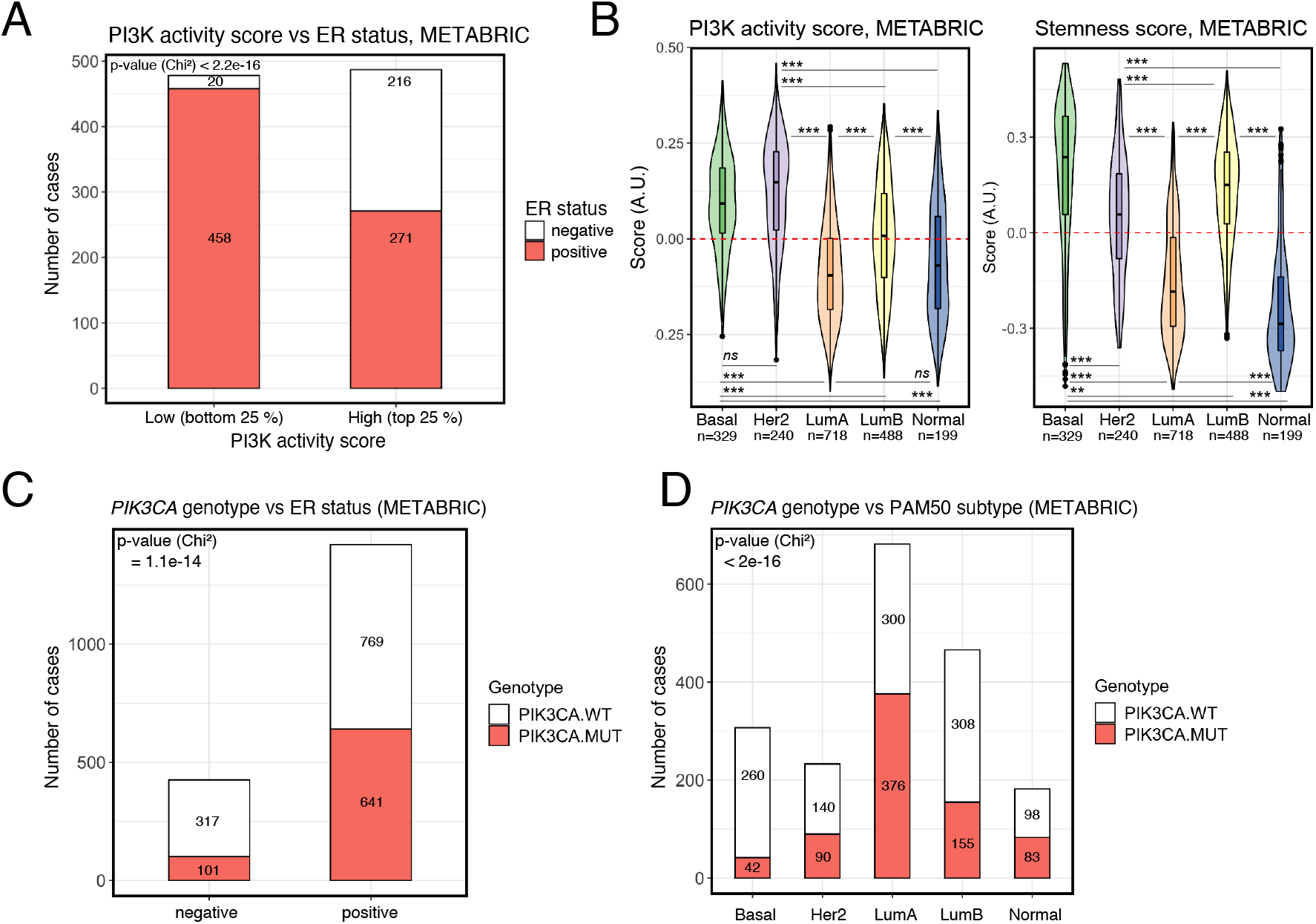
High PI3K activity and stemness scores, but not *PIK3CA* mutations, are enriched in aggressive breast cancer subtypes. **(A)** PI3K activity score distribution in METABRIC breast tumours stratified according to ER status. **(B)** PI3K activity and stemness score distributions across METABRIC breast cancers stratified according to PAM50 subtype; ** p ≤ 0.01, *** p ≤ 0.001 according to Tukey’s Honest Significant Differences method; *ns*: non-significant. **(C)** and **(D)** The distribution of *PIK3CA* wild-type (PIK3CA.WT) and mutant (PIK3CA.MUT) samples in METABRIC breast cancers, stratified according to ER status (C) or PAM50 subtype (D).

### PI3K and stemness scores, but not binary *PIK3CA* mutant status, predict prognosis in breast cancer

As expected, given the positive association between PI3K and stemness scores with tumour grade, both scores were negatively associated with patient survival in the METABRIC cohort, with a clear dosage relationship between the assessed scores and survival, including progressively worsened survival in tumours with high *vs* intermediate *vs* low scores (**Fig. 3A, 3B**). This relationship was not simply driven by the above-mentioned enrichment of high PI3K and stemness scores in more aggressive ER-negative tumours, as the prognostic power of both scores remained when evaluated in ER-positive tumours only (**Fig. 3C, 3D**). In contrast, although overall ER-negative cases with available survival data were limited in number, we in fact noticed a loss of prognostic power when evaluating the two scores in this breast cancer subset (**Fig. S1B, S1C**). Due to limited data, extensive survival analyses were not possible in TCGA breast cancers, however the negative association between PI3K activity “strength” and pan-breast cancer survival was reproduced (**Fig. S1D**).

**Fig. 3.**
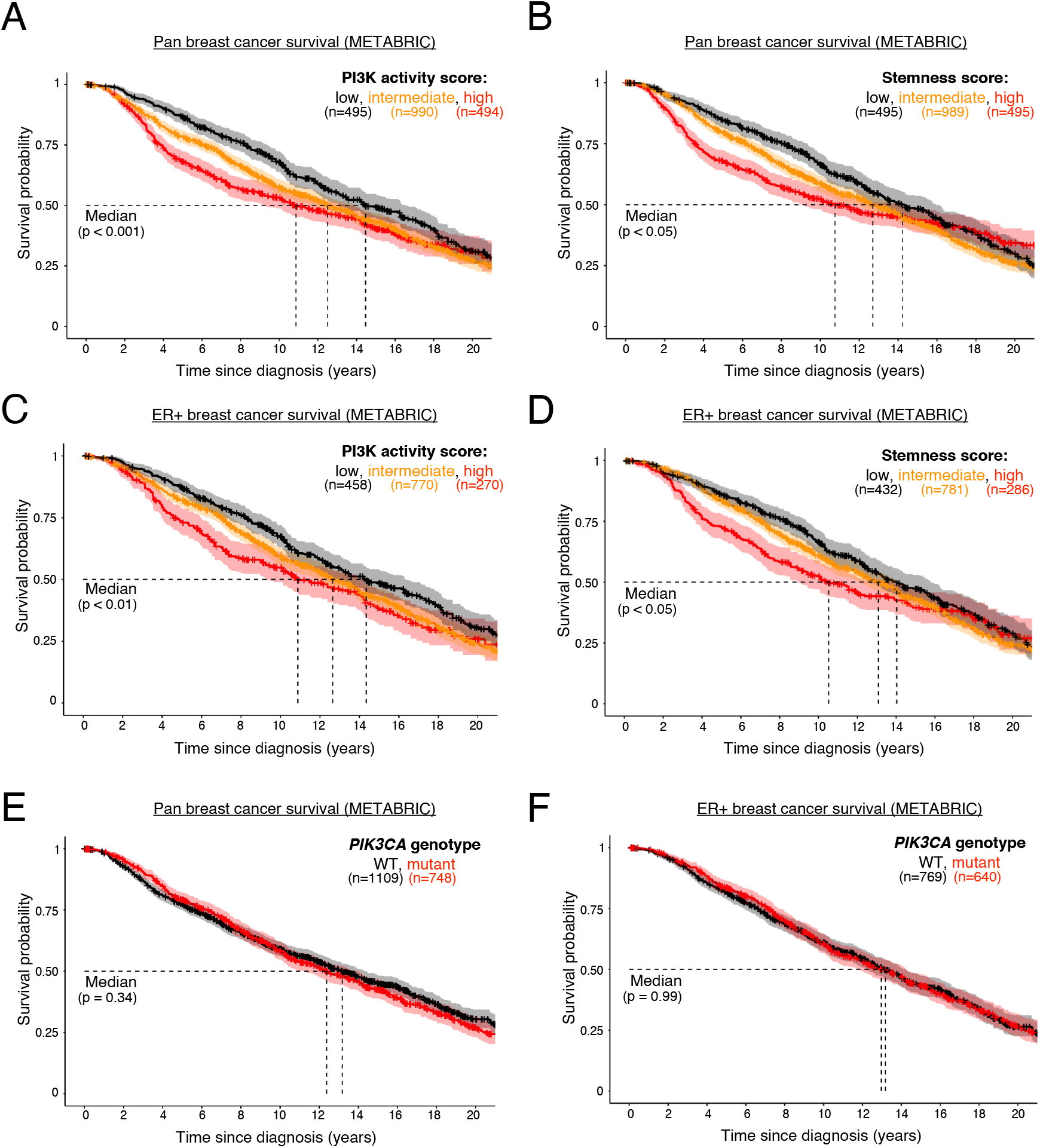
PI3K activity and stemness scores, but not *PIK3CA* genotype, are prognostic in ER+ breast cancer. Pan-breast cancer patient survival in METABRIC, as a function of PI3K activity **(A)** or stemness **(B)** score. Survival analysis in estrogen receptor (ER)-positive breast cancer patients, as a function of PI3K activity **(C)** or stemness **(D)** score. Low, intermediate and high classifications represent the bottom quartile, the interquartile range and the top quartile of the respective scores. **(E)** and **(D)** represent pan- and ER-positive breast cancer patient (METABRIC) survival, respectively, as a function of binary *PIK3CA* genotype. The mutant genotype captures only cases with activating missense mutations. The sample size for each panel and subgroup is indicated, and p-values were calculated using a log-rank test. The 95% confidence intervals are indicated by shading.

As previously reported [33–35], activating *PIK3CA* mutations had no prognostic power in pan-breast or ER-positive METABRIC tumours, despite their enrichment in the ER-positive cohort (**Fig. 3E, 3F**). Interestingly, however, the presence of *PIK3CA* mutations in ER-negative tumours appeared to be associated with worse prognosis (**Fig. S1E**).

### Stratification of breast cancers by mutant *PIK3CA* allele dosage reveals a biphasic relationship with PI3K activity and stemness scores

Given the divergent correlations between PI3K signalling scores and *PIK3CA* mutant status in the survival analyses, we next assessed the relationship between stemness/PI3K signalling scores and *PIK3CA* genotype, taking into account available information on mutant *PIK3CA* allele dosage on the basis of our previous work with TCGA tumours [8]. For METABRIC, we inferred *PIK3CA* copy number changes based on available information on allele gain/amplification in cBioPortal. For both cohorts, we specifically focused on tumours harbouring one or more hotspot *PIK3CA* alleles, given the well-established increased cellular activity of these mutants and their association with disease severity [36–39].

As PI3K pathway activation and tumour dedifferentiation can be triggered by a range of oncogenic hits, the relatively high PI3K and stemness scores in *PIK3CA*-WT breast cancers was not entirely surprising (**Fig. 4A, 4B**). It was, however, counterintuitive that the presence of a single oncogenic *PIK3CA* missense variant was associated with a substantial reduction in the stemness score and only a modest reduction in the PI3K score (**Fig. 4A, 4B**). Relative to tumours with a single *PIK3CA* mutant copy, those with multiple oncogenic *PIK3CA* copies exhibited higher PI3K and stemness scores (**Fig. 4A, 4B**). This relationship was lost upon simple binary classification based on *PIK3CA* genotypes (i.e. wild-type *vs* mutant) (**Fig. 4A, 4B**). The observed biphasic relationship also remained upon stratification of tumours according to genome doubling (data only available for TCGA samples; **Fig. 4C**).

**Fig. 4.**
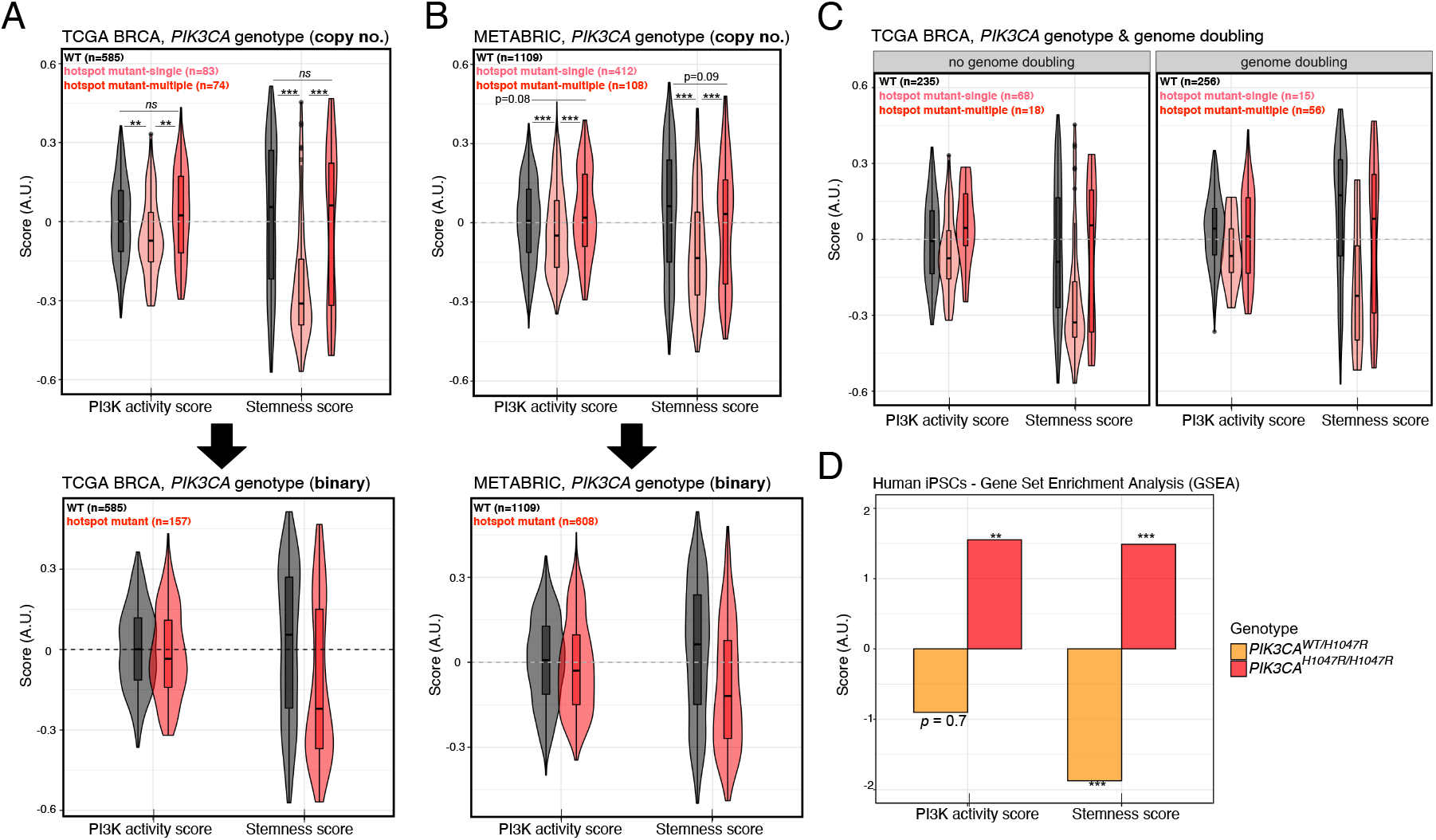
The presence of a single-copy, but not multi-copy, hotspot *PIK3CA* mutation is associated with lower PI3K activity and stemness score. **(A)** PI3K activity and stemness score distributions across TCGA breast cancers following stratification according to the presence or absence of single *vs* multiple copies of *PIK3CA* “hotspot” variants (C420R, E542K, E545K, H1047L, H1047R); ** p < 0.01, *** p < 0.001 according to one-way ANOVA with Tukey’s Honest Significant Differences method. **(B)** As in (A) but performed using METABRIC breast cancer transcriptomic and genomic data. **(C)** As in (A) but further stratified according to available genome doubling information. **(D)** Complementary GSEA-based PI3K and stemness score calculations using publicly-available transcriptomic data from iPSCs with heterozygous or homozygous *PIK3CA^H1047R^* expression [40]; enrichments are calculated relative to isogenic wild-type controls. ** p < 0.01, *** p < 0.001 for individual enrichments, according to FDR = 0.05 (Benjamini-Hochberg correction for multiple comparisons).

Surprised by this observation, we next asked whether the biphasic relationship between *PIK3CA* genotype and transcriptionally-derived PI3K/stemness scores could be recapitulated in a controlled cellular model. We turned to human induced pluripotent stem cells (iPSCs) that we engineered previously to harbour heterozygous or homozygous *PIK3CA^H1047R^* alleles, the only reported cellular models of heterozygous and homozygous *PIK3CA^H1047R^* expression on an isogenic background to date [40]. Using our previously reported high-depth transcriptomic data on *PIK3CA^WT/H1047R^* and *PIK3CA^H1047R/H1047R^* iPSCs [40], we performed conventional gene set enrichment analysis (GSEA) with the two gene set signatures used for PI3K and stemness score calculations in the breast cancer setting (MSigDB “HALLMARK_PI3K_AKT_MTOR_SIGNALING” and PluriNet, respectively). In line with their established biochemical and cellular phenotypes [8,40], homozygous *PIK3CA^H1047R^* iPSCs showed strong positive enrichment for both PI3K and stemness gene signatures (**Fig. 4D**). In contrast, their heterozygous *PIK3CA^H1047R^* counterparts presented with a strong negative enrichment for stemness, and no significant enrichment for the transcriptional PI3K signature (**Fig. 4D**). These patterns mirror those observed in human breast cancers and corroborate the existence of a previously unappreciated biphasic relationship between *PIK3CA* allele dosage and stemness.

### Stemness and PI3K activity scores are positively associated with proliferative and metabolic processes

Given the high depth and large sample size of the available breast cancer transcriptomic data, we next undertook a global analysis encompassing all 50 “hallmark” MSigDB gene sets and the PluriNet signature to identify relevant biological processes associated with breast cancer stemness and a high PI3K activity score. Such processes can be used to guide future experimental studies aimed at dissecting the molecular underpinnings of the observed relationships. To identify such associations, we applied GSVA to METABRIC and TCGA data to generate a score for each gene signature, followed by correlation analysis with hierarchical clustering. This global approach also allowed us to confirm that we are able to identify biologically-relevant gene signature clusters more broadly. For example, gene signatures associated with inflammatory processes clustered together according to strong pairwise positive correlations in both METABRIC and TCGA datasets (**Fig. 5A bottom cluster, Fig. 5B top left cluster**).

**Fig. 5.**
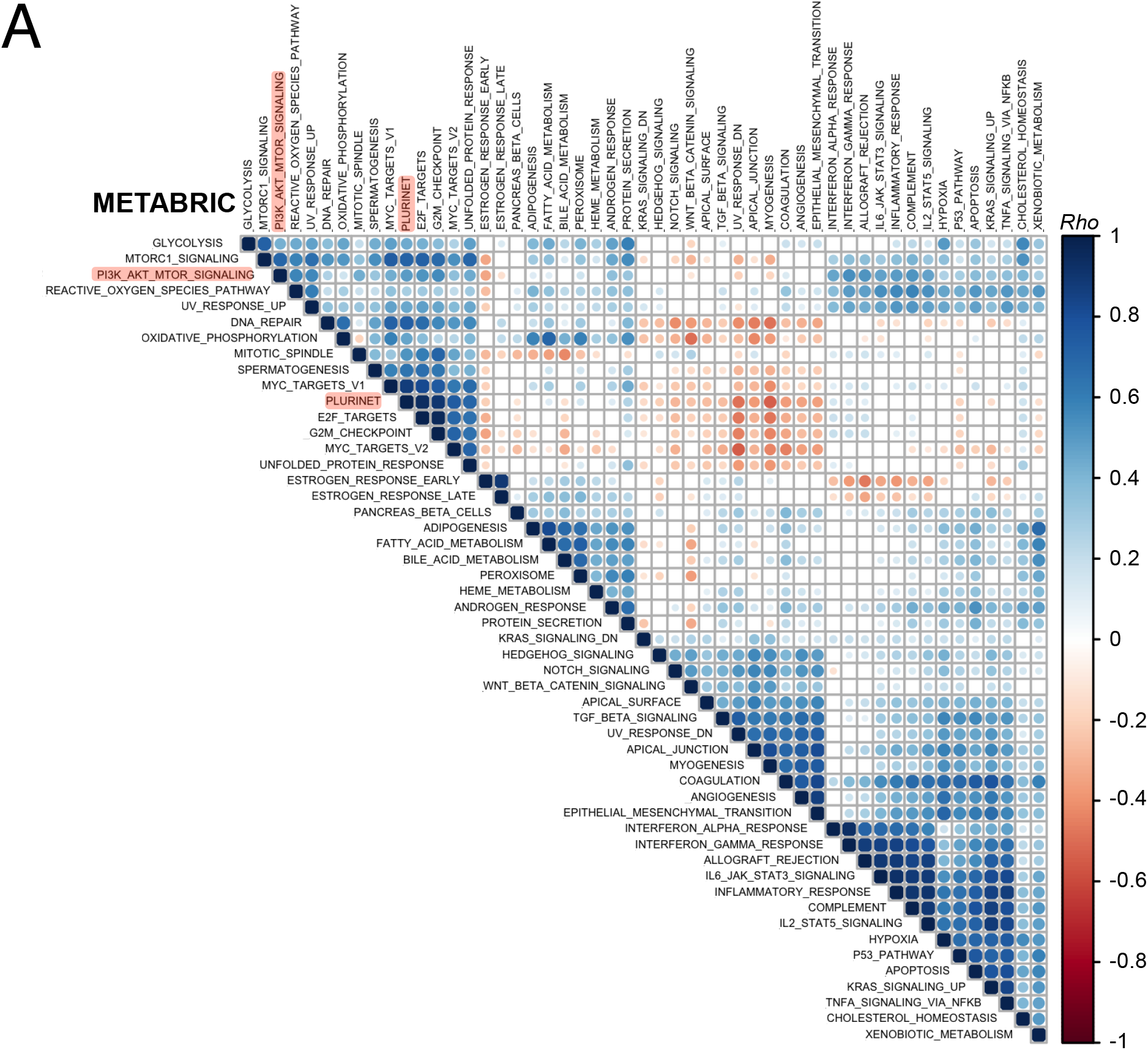
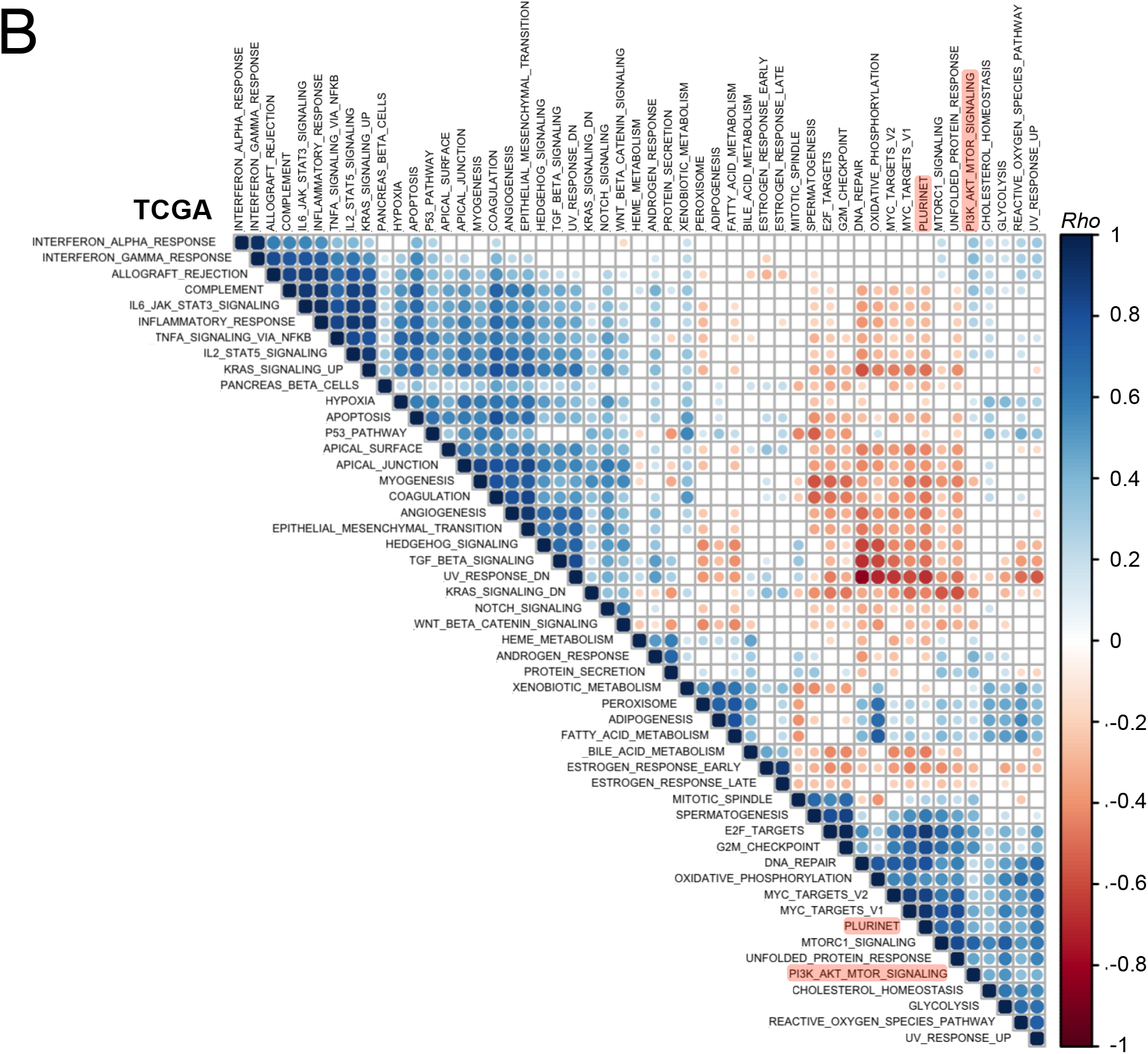
Breast cancer PI3K and stemness scores form a common cluster with proliferative and metabolic processes. Rank-based correlation analyses across METABRIC **(A)** and TCGA **(B)** GSVA-derived gene set enrichment scores, evaluating all 50 mSigDb Hallmark Gene Sets and PluriNet. Individual *Rho* coefficients are shown within the respective circles whose sizes are matched accordingly. Only significant correlations are shown (family-wise error rate < 0.05). The clusters were generated using unsupervised hierarchical clustering. The positions of PluriNet (stemness) and PI3K_AKT_MTOR (PI3K activity) signatures are highlighted in red.

Data from either cohort revealed a characteristic clustering pattern for PI3K and stemness scores, including strong positive associations with proliferative (e.g., “G2M_checkpoint”, “E2F_targets”, “MYC_targets”) and metabolic (e.g., “Glycolysis”, “Oxidative_phosphorylation”, “Reactive_oxygen_species”) gene signatures (**Fig. 5A, Fig. 5B**). These signatures shared few genes (**Fig. S1F**), ruling out technical artefacts as a source of the positive associations. Notably, the separate mTORC1 gene signature exhibited a much stronger correlation (Spearman’s rho = 0.7) with the stemness score compared with the PI3K_AKT_mTOR signature used to derive the PI3K activity score. Given a similarly high correlation between the PI3K_AKT_mTOR and mTORC1 signature scores (Spearman’s rho = 0.7), these data suggest that the observed relationship between PI3K and stemness in breast cancer may be driven by mTORC1-dependent processes.

## DISCUSSION

This study provides a comprehensive analysis of the relationship between PI3K signalling and stemness (or tumour dedifferentiation) using two large breast cancer transcriptomic datasets encompassing almost 3,000 primary tumours. We demonstrate a strong, positive relationship between transcriptionally-inferred PI3K pathway activity, stemness gene expression and histopathological tumour dedifferentiation. Importantly, we show that stratification of breast tumours according to single *vs* multiple copies of *PIK3CA* hotspot mutations results in distinct and near-opposite distributions with respect to PI3K signalling and stemness scores, an observation that is recapitulated in a controlled cell model system.

The PI3Kα-specific inhibitor alpelisib (Piqray/NVP-BYL719; Novartis) recently received approval for use in combination with the ER-antagonist fulvestrant in the treatment of ER-positive breast cancers. The benefit of this treatment was most notable in *PIK3CA*-mutant tumours, yet the predictive value of binary mutant classification was incomplete [5]. This is a common observation for single gene biomarkers in cancer and has long spurred discussions about the utility of phenotypic pathway signatures for clinical response prediction [41]. It is therefore interesting to note that while *PIK3CA* mutations are enriched in the ER-positive breast cancer subgroup, on average these tumours also feature lower PI3K signalling and stemness scores as inferred from our transcriptional footprint analyses. The opposite is true for ER-negative tumours. Given that the MSigDB hallmark PI3K_AKT_mTOR signature used in our study also encompasses mTORC1-related processes, in line with a strong correlation with the separate hallmark mTORC1 signature, our findings support a previous study reporting a negative relationship between the presence of a *PIK3CA* mutation and mTORC1 signalling in ER-positive/HER2-negative breast cancers [35]. As we show, however, simple binary classification of tumours into *PIK3CA* wild-type and mutant genotypes, without allele dosage considerations, is likely to have masked a more complex biological relationship. On the other hand, our study does not distinguish between AKT- and mTORC1-specific processes, which may nevertheless be important to consider for further mechanistic understanding and patient stratification [35,42,43].

Disentangling the apparent biphasic relationship between single *versus* multiple copies of *PIK3CA* mutation and stemness scores will require direct experimentation, but is likely to reflect context-dependent feedback loops within the intracellular signalling networks. Such feedback loops can result in non-intuitive and discontinuous outcomes upon different levels of activation of the same pathway, as demonstrated in our isogenic iPSC system with heterozygous and homozygous *PIK3CA^H1047R^* expression [8,40]. In general, our observations caution against the use of a binary *PIK3CA*-mutant-centric approach to predict PI3K pathway activity outcomes. Moreover, we note that numerous alternative genetic changes – including *PIK3CA* amplification, loss of *PTEN* or *INPP4B* – may converge on increased, and perhaps dose-dependent, PI3K pathway activation [3,30,32,44]. Importantly, such *PIK3CA* mutant-independent pathway activation is captured by the transcriptional footprint-based PI3K activity scores used in our study and will thus contribute to the values observed in non-*PIK3CA* mutant tumours.

While PI3K and stemness scores exhibit a strength-dependent negative association with patient survival pan-breast cancer as well as in ER-positive tumours, this prognostic power is not observed with binary genotype-based *PIK3CA* classification. Paradoxically, however, *PIK3CA* mutations have prognostic power in ER-negative tumours, in contrast to PI3K signalling and stemness scores. This raises the question whether subgroups defined by differences in *PIK3CA* mutant status and PI3K signalling/stemness scores differ in their response to PI3Kα-targeted therapy.

It is also notable that our correlation analyses of breast cancer transcriptomes identified a PI3K/stemness cluster encompassing key processes associated with the MYC regulatory module in pluripotent stem cells [45]; a module previously shown to be active in various cancers and predictive of cancer outcome [46]. Moreover, computational analyses of iPSCs with homozygous *PIK3CA^H1047R^* expression identified MYC as a central hub connecting the PI3K, TGFβ and pluripotency networks in these cells [40]. Recently, *PIK3CA^H1047R^/KRAS^G12V^* double knock-in breast epithelial cells were also shown to exhibit a high MYC transcriptional signature, when compared to single-mutant counterparts [47]. Collectively, the recurrent appearance of MYC in these independent analyses raises the possibility that this transcription factor governs the mechanistic link between stemness and PI3K signalling strength in pluripotent stem cells and breast cancer. Experimental studies will be required to test this hypothesis, alongside a potential involvement of mTORC1 as suggested by the observed strong positive correlation between the mTORC1 signature and stemness/MYC signatures.

A limitation of the current and previous bulk-tissue transcriptomic analyses is that they cannot determine (1) whether the observed correlations reflect mechanistic links or spurious associations caused by a confounder variable that influences two or more processes independently and (2) to what extent the observed transcriptomic scores are driven by changes in the subcellular composition, tumour cell type-specific phenotypic alterations, and/or non-cell-autonomous interactions with the stroma. Nevertheless, given our ability to reproduce key observations in a controlled cell model system, our analyses of the relationship between PI3K signalling dose and stemness in breast cancer may prove useful in guiding future experimental studies aimed at identifying the exact molecular underpinnings. Since we know that heterozygous *PIK3CA^H1047R^* iPSCs exhibit moderate PI3K pathway activation at the biochemical level [8,40], the fact that this is not captured in a positive transcriptional footprint-based PI3K score is worth noting. Combined with the observation of an apparent decrease in the PI3K score in tumours with a single copy of a hotspot *PIK3CA* mutation, we surmise that this may reflect feedback mechanisms that limit the influence of intermediate PI3K pathway activation but that are not sufficient in the face of stronger activity. This warrants further studies as it may have important consequences for targeting of tumours with a high *versus* low transcriptionally-inferred PI3K score. It is also worth noting that previous protein-based signalling studies of breast cancer cell lines and tumours with and without *PIK3CA* mutations found that *PIK3CA* mutations were associated with lower and/or inconsistent PI3K pathway activation [30,33,35].

Finally, on the basis of the presented analyses, it will be of interest to evaluate the predictive power of a combined assessment of *PIK3CA* genotype and phenotypic PI3K/stemness scores in patient stratification for clinical trials with PI3K pathway inhibitors and, given the well-established implication of PI3K signalling in therapeutic response and resistance, with other cancer therapies.

## MATERIALS AND METHODS

### Data and materials availability

All computational analyses were conducted in R [48]. The below represent summaries of the applied methods. Detailed step-by-step workflows on each breast cancer cohort, can be found on the accompanying Open Science Framework (OSF) page: https://osf.io/g8rf3/wiki/home/. This also contains all source datasets and key output data tables as well as figure. As indicated in the accompanying scripts, all relevant packages were sourced either from CRAN or Bioconductor (via BiocManager [49]). Figures were produced using the *ggplot2* package [50].

Further information requests should be directed to and will be fulfilled by the corresponding authors, Ralitsa R. Madsen and Bart Vanhaesebroeck.

### METABRIC and TCGA data access and pre-processing

Normalised microarray-based gene expression for METABRIC breast tumour samples were obtained from Curtis et al. [51], and clinical data from Rueda et al. [52]. The relevant METABRIC mutation data were downloaded from cBioPortal in January (mutation-only) and March (mutation and copy number) 2020 [53]. TCGA breast invasive carcinoma (BRCA) RNAseq, mutational and clinical data were retrieved from the GDC server (legacy database) using the *TCGAbiolinks* package [54], with additional mutation data retrieved from cBioPortal in January 2021 (for exact details, see the OSF-deposited RNotebooks). The *TCGAbiolinks* package was also used for subsequent quantile filtering (quantile value = 0.4) of lowly-expressed gene and removal of tumour samples with low purity (cpe = 0.6). The resulting raw RSEM counts were normalised with the TMM method [55] and log2-transformed using the voom() function in the *limma* package prior to downstream use in GSVA computations. The TCGA BRCA mutation data with available copy number estimates for individual mutations were obtained from Madsen et al. [8] and merged with the mutation data from cBioPortal. Multiple allele copies were defined as those having mut.multi > 1.5.

To analyse the relationships between *PIK3CA* genotype and PI3K/stemness scores, *PIK3CA* mutant datasets were subset for focus on hotspot *PIK3CA* variants only (C420R, E542K, E545K, H1047L, H1047R), excluding samples containing both a hotspot and a non-hotspot variant. The classification of hotspot vs non-hotspot variants was based on known clinical significance and frequency in patients with overgrowth caused by a single activating *PIK3CA* mutations [38]. Mutation data underwent manual checks to exclude samples with ambiguous genotype calls as well as all silent mutations.

### Calculation of transcription-based signature scores

The “HALLMARK_PI3K_AKT_MTOR_SIGNALING” and PluriNet gene sets were retrieved from The Molecular Signature Database (MSigDb) using the *msigdbr* package [56]. Note that the “HALLMARK_PI3K_AKT_MTOR_SIGNALING” gene set also includes mTORC1-dependent gene expression changes, in contrast to other studies which have sought to separate AKT- and mTORC1-driven gene expression changes [42,43]. Categorisation of scores into “low”, “intermediate” and “high” was based on the 0.25 quantile, the interquartile range, and the 0.75 quantile, respectively. The stemness signature used by Miranda *et al*. [22] was retrieved from the accompanying supplementary material. Individual scores for each of these signatures were computed with the *GSVA* package, using the default Gaussian kernel and ESdiff enrichment values as output [29].

The PROGENy package was used to obtain a PI3K score according to a linear model based on pathway-responsive genes as described in Ref. [23].

The TCGAnalyze_Stemness() function in *TCGAbiolinks* was used to calculate a stemness score according to the machine learning model-based mRNAsi signature reported by Malta et al. [21].

Transcriptomic data for human induced pluripotent stem cells with wild-type *PIK3CA* or heterozygous/homozygous *PIK3CA^H1047R^* were available from Ref. [40]. Fast gene set enrichment analysis (fgsea) [57] with the “HALLMARK_PI3K_AKT_MTOR_SIGNALING” and “PluriNet” gene sets were performed using the *t* statistic for all genes from comparisons between *PIK3CA^WT/H1047R^* (heterozygous) vs wild-type and *PIK3CA^H1047R/H1047R^* vs wild-type sampes. Multiple comparison adjustment were performed using the intrinsic *fgsea()* function settings (FDR = 0.05), except that nominal p-values were calculated from 100,000 permutations for increased stringency.

### Statistical analyses

Linear models were used to assess the significance of the relationship between stemness and PI3K scores in both METABRIC and TCGA breast cancer cohorts. One-way ANOVA followed by Tukey’s Honest Significant Differences (HSD) method was used to perform pairwise significance testing with multiple comparison adjustments (adjusted p-value < 0.05) when evaluating grade- and cancer subtype-specific differences in PI3K/stemness scores across the METABRIC cohort; similar analyses were not performed with the TCGA breast cancer data due to smaller sample size and incomplete grading information. ANOVA with Tukey’s HSD was also used to evaluate the significance of the relationships between *PIK3CA* genotype and PI3K/stemness scores across both cohorts. For linear models as well as ANOVAs, the residuals were examined to confirm that model assumptions were met. The only assumption that was violated was that of normality; however, given the large sample size, this violation is expected to have a minimal impact on model validity [58].

Differences in categorial PI3K/stemness score (“low”, “intermediate”, “high”) distributions across tumour subtypes and/or genotypes were assessed using a Chi-squared goodness-of-fit test. The relationship between PI3K/scores and survival was assessed using a non-parametric log-rank test.

Pairwise correlation analyses and hierarchical clustering of signature scores were performed using Spearman’s rank correlation and the Ward.D2 method (available through R package *corrplot*; https://github.com/taiyun/corrplot). The associated p-values were adjusted for multiple comparisons using the Bonferroni method (family-wise error rate < 0.05).

## Acknowledgements

We are grateful to Dr Neil Vasan (Memorial Sloan Kettering Cancer Center, New York) for excellent feedback on the manuscript. We would also like to thank the cancer community behind the TCGA/METABRIC datasets, as well as data scientists developing the above-mentioned analysis tools, for making them publicly available and thus enabling the completion of this study.

## Supplementary Material

### Other supplementary materials for this manuscript include the following

Separate source code file for analysis of METABRIC/TCGA PI3K activity and stemness scores (https://osf.io/g8rf3/wiki/home/ & doi: 10.17605/OSF.IO/G8RF3)
Supplementary Table 1: mSigDb “HALLMARK_PI3K_AKT_MTOR_SIGNALING” gene list
Supplementary Table 2: mSigDb “MUELLER_PLURINET” gene list

**Fig. S1.**
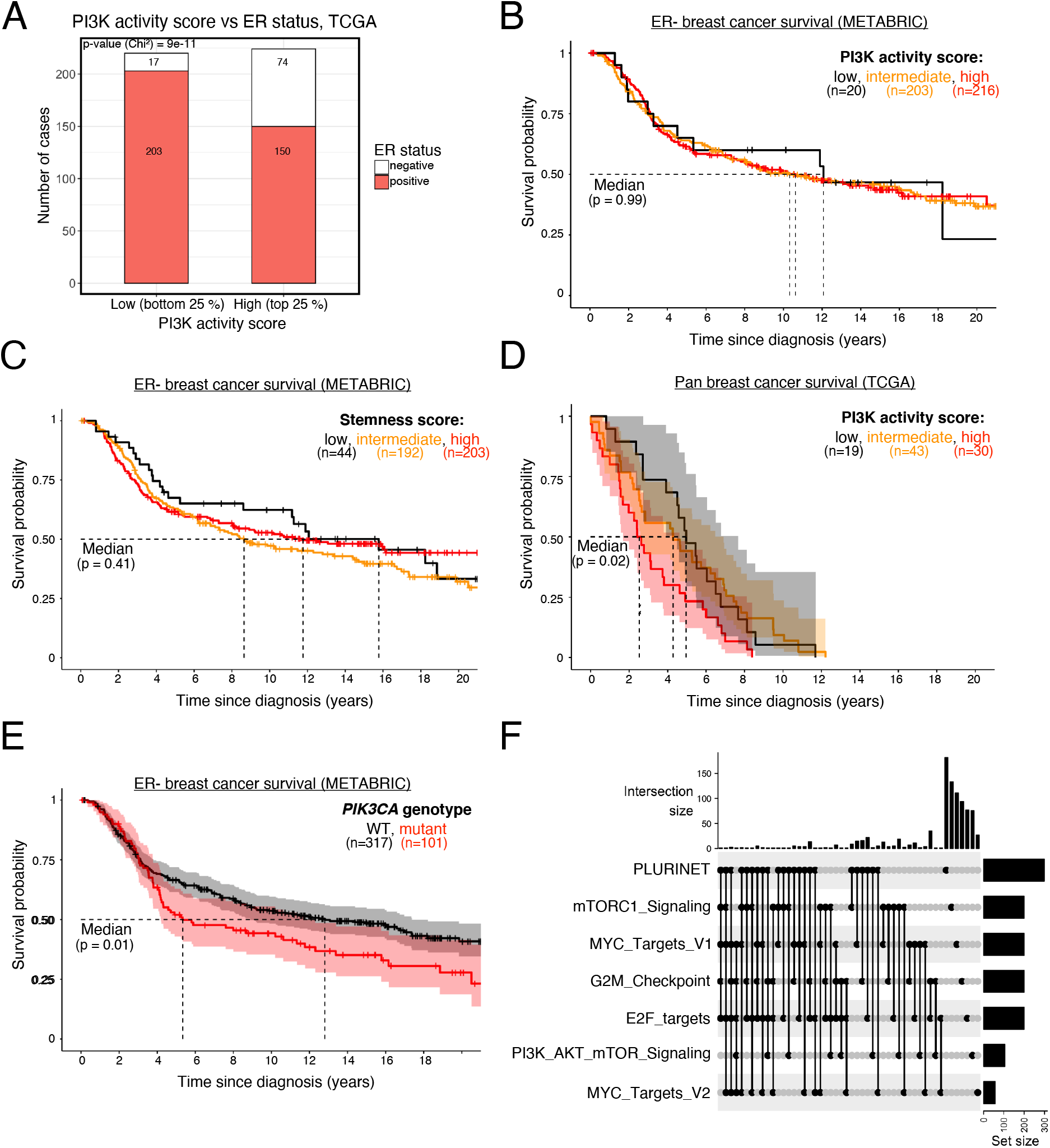
**(A)** PI3K activity score distribution in TCGA breast tumours stratified according to ER status. Survival analysis in estrogen receptor (ER)-negative breast cancer patients, as a function of PI3K activity **(B)** or stemness **(C)** score. **(D)** Pan-breast cancer patient survival in TCGA, as a function of PI3K activity score. **(E)** ER-negative breast cancer patient (METABRIC) survival as a function of binary *PIK3CA* genotype. The sample size for each panel and subgroup is indicated, and p-values were calculated using a log-rank test; where shown, the 95% confidence intervals are indicated by shading. **(F)** UpSet plot showing intersection set sizes across the specified gene set combinations.

## References

1 Chang, M. T. et al. (2015) Identifying recurrent mutations in cancer reveals widespread lineage diversity and mutational specificity. Nat. Biotechnol. 34, 155–163.

2 Campbell, P. J. et al. (2020) Pan-cancer analysis of whole genomes. Nature 578, 82–93.

3 Sanchez-Vega, F. et al. (2018) Oncogenic Signaling Pathways in The Cancer Genome Atlas. Cell 173, 321–337.e10.

4 Vanhaesebroeck, B. et al. (2021) PI3K inhibitors are finally coming of age. Nat. Rev. Drug Discov.

5 André, F. et al. (2019) Alpelisib for PIK3CA-mutated, hormone receptor-positive advanced breast cancer. N. Engl. J. Med. 380, 1929–1940.

6 Madsen, R. R. et al. (2018) Cancer-Associated PIK3CA Mutations in Overgrowth Disorders. Trends Mol. Med. 24, 856–870.

7 Yuan, T. L. and Cantley, L. C. (2008) PI3K pathway alterations in cancer: variations on a theme. Oncogene 27, 5497–5510.

8 Madsen, R. R. et al. (2019) Oncogenic PIK3CA promotes cellular stemness in an allele dose-dependent manner. Proc. Natl. Acad. Sci. 116, 8380–8389.

9 Vasan, N. et al. (2019) Double PIK3CA mutations in cis increase oncogenicity and sensitivity to PI3Kα inhibitors. Science. 366, 714–723.

10 Saito, Y. et al. (2020) Landscape and function of multiple mutations within individual oncogenes. Nature 582, 95–99.

11 Van Keymeulen, A. et al. (2015) Reactivation of multipotency by oncogenic PIK3CA induces breast tumour heterogeneity. Nature 525, 119–23.

12 Koren, S. et al. (2015) PIK3CA(H1047R) induces multipotency and multi-lineage mammary tumours. Nature 525, 114–8.

13 Hanker, A. B. et al. (2013) Mutant PIK3CA accelerates HER2-driven transgenic mammary tumors and induces resistance to combinations of anti-HER2 therapies. Proc. Natl. Acad. Sci. U. S. A. 110, 14372–14377.

14 van Veen, J. E. et al. (2019) Mutationally-activated PI3’-kinase-α promotes de-differentiation of lung tumors initiated by the BRAFV600E oncoprotein kinase. Elife 8, 1–33.

15 Riemer, P. et al. (2017) Oncogenic β-catenin and PIK3CA instruct network states and cancer phenotypes in intestinal organoids. J. Cell Biol. 216, 1567–1577.

16 Du, L. et al. (2016) Overexpression of PIK3CA in murine head and neck epithelium drives tumor invasion and metastasis through PDK1 and enhanced TGFβ signaling. Oncogene 35, 4641–4652.

17 Meyer, D. S. et al. (2011) Luminal expression of PIK3CA mutant H1047R in the mammary gland induces heterogeneous tumors. Cancer Res. 71, 4344–4351.

18 Intlekofer, A. M. and Finley, L. W. S. (2019) Metabolic signatures of cancer cells and stem cells. Nat. Metab. 1, 177–188.

19 Ben-Porath, I. et al. (2008) An embryonic stem cell-like gene expression signature in poorly differentiated aggressive human tumors. Nat. Genet. 40, 499–507.

20 Palmer, N. P. et al. (2012) A gene expression profile of stem cell pluripotentiality and differentiation is conserved across diverse solid and hematopoietic cancers. Genome Biol. 13.

21 Malta, T. M. et al. (2018) Machine Learning Identifies Stemness Features Associated with Oncogenic Dedifferentiation. Cell 173, 338–354.e15.

22 Miranda, A. et al. (2019) Cancer stemness, intratumoral heterogeneity, and immune response across cancers. Proc. Natl. Acad. Sci. U. S. A. 116, 9020–9029.

23 Schubert, M. et al. (2018) Perturbation-response genes reveal signaling footprints in cancer gene expression. Nat. Commun. 9.

24 Liberzon, A. et al. (2015) The Molecular Signatures Database Hallmark Gene Set Collection. Cell Syst. 1, 417–425.

25 Szalai, B. and Saez-Rodriguez, J. (2020) Why do pathway methods work better than they should? FEBS Lett. 594, 4189–4200.

26 Subramanian, A. et al. (2017) A Next Generation Connectivity Map: L1000 Platform and the First 1,000,000 Profiles. Cell 171, 1437–1452.e17.

27 Lamb, J. et al. (2006) The Connectivity Map: using gene-expression signatures to connect small molecules, genes, and disease. Science. 313, 1929–35.

28 Müller, F. J. et al. (2008) Regulatory networks define phenotypic classes of human stem cell lines. Nature 455, 401–405.

29 Hänzelmann, S. et al. (2013) GSVA: Gene set variation analysis for microarray and RNA-Seq data. BMC Bioinformatics 14.

30 López-Knowles, E. et al. (2010) PI3K pathway activation in breast cancer is associated with the basal-like phenotype and cancer-specific mortality. Int. J. Cancer 126, 1121–1131.

31 Creighton, C. J. et al. (2010) Proteomic and transcriptomic profiling reveals a link between the PI3K pathway and lower estrogen-receptor (ER) levels and activity in ER+ breast cancer. Breast Cancer Res. 12.

32 Koboldt, D. C. et al. (2012) Comprehensive molecular portraits of human breast tumours. Nature 490, 61–70.

33 Stemke-Hale, K. et al. (2008) An integrative genomic and proteomic analysis of PIK3CA, PTEN, and AKT mutations in breast cancer. Cancer Res. 68, 6084–6091.

34 Pérez-Tenorio, G. et al. (2007) PIK3CA mutations and PTEN loss correlate with similar prognostic factors and are not mutually exclusive in breast cancer. Clin. Cancer Res. 13, 3577–3584.

35 Loi, S. et al. (2010) PIK3CA mutations associated with gene signature of low mTORC1 signaling and better outcomes in estrogen receptor – positive breast cancer. Proc. Natl. Acad. Sci. U. S. A. 107, 10208–10213.

36 Dogruluk, T. et al. (2015) Identification of Variant-Specific Functions of PIK3CA by Rapid Phenotyping of Rare Mutations. Cancer Res. 75, 5341–54.

37 Zhang, Y. et al. (2017) A Pan-Cancer Proteogenomic Atlas of PI3K/AKT/mTOR Pathway Alterations. Cancer Cell 31, 820–832.e3.

38 Mirzaa, G. et al. (2016) PIK3CA-associated developmental disorders exhibit distinct classes of mutations with variable expression and tissue distribution. JCI Insight 1, 1–18.

39 Kuentz, P. et al. (2017) Molecular diagnosis of PIK3CA-related overgrowth spectrum (PROS) in 162 patients and recommendations for genetic testing. Genet. Med. 19, 989–997.

40 Madsen, R. R. et al. (2021) NODAL/TGFβ signalling mediates the self-sustained stemness induced by PIK3CA H1047R homozygosity in pluripotent stem cells. Dis. Model. Mech. dmm. 048298.

41 Watters, J. W. and Huang, P. S. (2009) Can phenotypic pathway signatures improve the prediction of response to PI3K pathway inhibitors? Drug Discov. Today Ther. Strateg. 6, 57–62.

42 Sonnenblick, A. et al. (2019) pAKT pathway activation is associated with PIK3CA mutations and good prognosis in luminal breast cancer in contrast to p-mTOR pathway activation. Breast Cancer 5, 1–9.

43 Creighton, C. J. (2007) A gene transcription signature of the Akt/mTOR pathway in clinical breast tumors. Oncogene 26, 4648–4655.

44 Liu, H. et al. (2020) The INPP4B Tumor Suppressor Modulates EGFR Trafficking and Promotes Triple Negative Breast Cancer. Cancer Discov. CD-19–1262.

45 Fagnocchi, L. and Zippo, A. (2017) Multiple Roles of MYC in Integrating Regulatory Networks of Pluripotent Stem Cells. Front. Cell Dev. Biol. 5, 1–19.

46 Kim, J. et al. (2010) A Myc Network Accounts for Similarities between Embryonic Stem and Cancer Cell Transcription Programs. Cell 143, 313–324.

47 Bell, C. M. et al. (2019) PIK3CA Cooperates with KRAS to Promote MYC Activity and Tumorigenesis via the Bromodomain Protein BRD9. Cancers (Basel). 11, 1634.

48 R Core Team. (2019) R: A language and environment for statistical computing. R Found. Stat. Comput.

49 Morgan, M. (2019) BiocManager: Access the Bioconductor Project Package Repository.

50 Wickham, H. (2016) ggplot2: Elegant Graphics for Data Analysis.

51 Curtis, C. et al. (2012) The genomic and transcriptomic architecture of 2,000 breast tumours reveals novel subgroups. Nature 486, 346–352.

52 Rueda, O. M. et al. (2019) Dynamics of breast-cancer relapse reveal late-recurring ERpositive genomic subgroups. Nature 567, 399–404.

53 Cerami, E. et al. (2012) The cBio Cancer Genomics Portal: An open platform for exploring multidimensional cancer genomics data. Cancer Discov. 2, 401–404.

54 Colaprico, A. et al. (2016) TCGAbiolinks: An R/Bioconductor package for integrative analysis of TCGA data. Nucleic Acids Res. 44, e71.

55 Robinson, M. D. and Oshlack, A. (2010) A scaling normalization method for differential expression analysis of RNA-seq data. Genome Biol. 11, R25.

56 Dolgalev, I. (2020) msigdbr: MSigDB Gene Sets for Multiple Organisms in a Tidy Data Format.

57 Korotkevich, G. et al. (2016) Fast gene set enrichment analysis. bioRxiv 1-29.

58 Schmidt, A. F. and Finan, C. (2018) Linear regression and the normality assumption. J. Clin. Epidemiol. 98, 146–151.

